# Monoterpenes alter TAR1-driven physiology in *Drosophila* species

**DOI:** 10.1101/2020.06.26.173732

**Authors:** Luca Finetti, Lasse Tiedemann, Xiaoying Zhang, Stefano Civolani, Giovanni Bernacchia, Thomas Roeder

**Affiliations:** Department of Life Sciences and Biotechnology, University of Ferrara, Ferrara, Italy; Laboratory of Molecular Physiology, Department of Zoology, Kiel University, Kiel, Germany; InnovaRicerca s.r.l. Monestirolo, Ferrara, Italy; German Center for Lung Research (DZL), Airway Research Center North (ARCN), Kiel, Germany

**Keywords:** *Drosophila*, Monoterpenes, Tyramine receptor, Metabolism, Behaviour

## Abstract

Monoterpenes are molecules with insecticide properties whose mechanism of action is however not completely elucidated. Furthermore, they seem to be able to modulate the monoaminergic system and several behavioural aspects in insects. In particular, tyramine (TA) and octopamine (OA) and their associated receptors orchestrate physiological processes such as feeding, locomotion and metabolism. Here we show that monoterpenes not only act as biopesticides in *Drosophila* species but can cause complex behavioural alterations that require a functional type 1 tyramine receptors (TAR1s). Variations in metabolic traits as well as locomotory activity were evaluated in both *Drosophila suzukii* and *Drosophila melanogaster* after treatment with three monoterpenes. A TAR1^−/−^ *D. melanogaster* strain was used to better understand the relationships between the receptor and monoterpenes-related behavioural changes. Immunohistochemistry analysis revealed that, in the *D. melanogaster* brain, TAR1 appeared to be expressed in areas controlling metabolism. In comparison to the *D. melanogaster* wild type, the TAR^−/−^ flies showed a phenotype characterized by higher triglyceride levels and food intake as well as lower locomotory activity. The monoterpenes, tested at sublethal concentrations, were able to induce a downregulation of the TAR1 coding gene in both *Drosophila* species. Furthermore, monoterpenes also altered the behaviour in *D. suzukii* and *D. melanogaster* wild types 24 h after a continuous monoterpene exposure. Interestingly, they were ineffective in modifying the physiological performances of TAR1^−/−^ flies. In conclusion, it appears that monoterpenes not only act as biopesticides for *Drosophila* but they can also interfere with its behaviour and metabolism in a TAR1-dependent fashion.

## Introduction

*Drosophila suzukii* Matsumura (Diptera: Drosophilidae), commonly known as “Spotted Wing Drosophila”, is one of the few Drosophilidae that can lay its eggs on healthy fruits before they becomes fully ripe (Walsh et al., 2011; Lee et al., 2011). *D. suzukii* is able to infest most of the fruit and vine species worldwide with a particular preference for small fruits (Rota-Stabelli et al., 2013). This species causes serious damages to the horticultural economy especially in South-East Asia and its presence has been recently reported also in North America and Europe (Asplen et al., 2015). Moreover, *D. suzukii* can spread rapidly (seven to fifteen generations – year) and has a remarkable ability to adapt to different climatic conditions and host plants (Cini et al., 2012). Chemical pesticides are the main *D. suzukii* control agents, but they need frequent enforcements due to the numerous generations that occur during one crop season. Nonetheless, repetitive treatments may increase resistance development and have a negative impact on beneficial insects (Desneux et al., 2007; Haviland & Beers, 2012). Alternative and more sustainable control strategies are constantly under investigation (Schetelig et al., 2017). Currently, research on the biology, genetics, as well as physiology of *D. suzukii* has gained interest in order to develop new tools for a more effective and environmentally sensitive pest management. Essential oils (EOs) as botanical pesticides are among the most promising pest control methods for future applications. In fact, studies performed in the last decade showed that pesticides based on plant essential oils and their constituents (terpenes) are effective against a large number of insects (Bakkali et al., 2008; Isman, 2020). Members of the Drosophilidae family, *D. suzukii* included, are particularly sensitive to EO based pesticides (Park et al., 2016, Kim et al., 2016; Zhang et al., 2016; Dam et al., 2019). Most of EOs are complex mixtures of two predominant classes of molecules, terpenes and phenylpropanoids (Regnault-Roger et al., 2012). Although it is clear that EOs have toxic effects against pest insects, their mechanism of action is still unclear (Blenau et al., 2011; Jankowska et al., 2018). Typically, they are able to reduce or disrupt insect growth at several life stages (Konstantopoulou et al., 1992). It has been shown that terpenes can interact with P450 cytochromes, which are involved in insecticide detoxification processes (Jensen et al., 2006; Liao et al., 2016). Some monoterpenes, for example thymol, may induce neuronal degeneration through a direct interaction with GABA receptors (Priestley et al., 2003) or via acetylcholinesterase inhibition (Houghton et al., 2006; Park et al., 2016). Moreover, monoterpenes might interact with the octopamine/tyramine system, analogous to the adrenergic system present in the vertebrates (Enan, 2001; Kostyukovsky et al., 2002; Enan, 2005a; Enan, 2005b; Price & Berry, 2006; Gross et al., 2017; Finetti et al., 2020).

In insects, the main biogenic amines are dopamine (DA), serotonin (5-HT), octopamine (OA) and tyramine (TA). Together, they control and modulate a broad range of biological functions essential for the insects life (Roeder et al., 2003). The insect’s nervous system contains high levels of OA and TA, suggesting a role as neurotransmitters (Ohta & Ozoe, 2014), but also as neuromodulators and neurohormones in a wide variety of physiological processes (Pauls et al., 2018).

Originally, TA was considered only as an intermediate product necessary for the synthesis of OA. Nevertheless, today it is known that TA and OA perform important functions independently of each other (Roeder, 2005; Lange, 2009; Roeder, 2020). TA triggers its physiological effects by interacting with and activating the corresponding receptors, belonging to the G Protein-Coupled Receptors (GPCR) family (Evans & Maqueira, 2005). Tyramine receptors (TARs) play important roles in modulating the biology, physiology and behaviour of invertebrates (Ohta & Ozoe, 2014). In fact, either the inhibition or the over stimulation of TARs can lead to the death of the insect as well as interfere with physical fitness and reproductive capacity (Audsley & Down, 2015). These receptors are classified into two main groups based on their structure and activity: tyramine receptors type 1 (TA/OA or TAR1) on one hand and tyramine receptors type 2 and 3 on the other (TAR2 and TAR3) (Wu et al., 2014). TAR1 transcripts localization analysis provides clues to understand its physiological roles. In *D. melanogaster*, the receptor is highly expressed in the central nervous system CNS (Saudou et al., 1990; El-Kholy et al., 2015). A similar expression pattern has been observed also in *D. suzukii, R. prolixus, C. suppressalis, P. xylostella, M. brassicae* and *A. ipsilon* suggesting a crucial role for TA as neuromodulator and neurotransmitter (Wu et al., 2013; Hana & Lange, 2017; Ma et al., 2019; Brigaud et al., 2009; Duportets et al., 2010; Finetti et al., 2020). Several studies have reported the importance of TA, through its interaction with TARs, in a variety of processes including olfaction, reproduction, flight, locomotion and metabolic traits (Lange, 2009; Neckameyer & Leal, 2017; Roeder, 2020). In particular, TA appears to play a role in locomotor modulation (Saraswati et al., 2004; Hardie et al., 2007; Rillich et al., 2013; Schützler et al., 2019), in egg-laying behaviour (Donini & Lange, 2004; Fuchs et al., 2014), in sex pheromone production (Hirashima et al., 2007), in metabolic traits including the regulation of energy expenditure (Brembs et al., 2007) and hormone release (Roeder, 2020). Despite the physiological importance of TA in invertebrates, little is known about tyramine receptors. In 2000 Kutsukake and co-workers characterized *D. melanogaster hono*, a mutant line with an impaired TAR1, exhibiting a different behaviour towards repellent odours. Furthermore, Li et al. (2017) have showed that TAR1 deficient flies exhibit significant changes in the metabolic control such as higher body fat, lower starvation resistance and movement activity. Similar TAR1-mediated metabolic alterations were observed by Ishida & Ozaki (2011) in starved flies. Nevertheless, the existence of a crosstalk between the tyraminergic system and other systems, such as the octopaminergic and dopaminergic, makes it difficult to precisely dissect the physiological processes controlled by TA (Li et al., 2016).

In the last few years, several studies have suggested that TAR1 might be an interesting target for insecticides, specifically for bioinsecticides. For example, monoterpenes appear to be able to interact with TAR1 directly. In particular, Enan (2005b) was the first to describe an agonistic effect of several monoterpenes (thymol, carvacrol, α-terpineol and eugenol) on *D. melanogaster* TAR1. However, the same monoterpenes did not show this pharmacological profile on *D. suzukii* and *Rhipicephalus microplus* TAR1 receptors. They acted instead as positive allosteric modulators, increasing the potency of TA activity (Gross et al., 2017; Finetti et al., 2020). Furthermore, a recent study from our lab has described a possible molecular mechanism underlying the toxicity of these molecules towards insects (Finetti et al., 2020). In particular, the observed downregulation of *D. suzukii* TAR1 (DsTAR1) after monoterpene exposure might represent a compensatory mechanism in response to the enhanced receptor signalling due to the positive allosteric modulatory effect of monoterpenes on the receptor.

The current study presents a detailed investigation on *D. suzukii* behaviour upon monoterpenes treatment, in order to understand whether the *DsTAR1* downregulation could affect fitness and physiology. Furthermore, a *D. melanogaster* mutant line impaired in TAR1 was used as a control to compare the effects of chronic TAR1 absence on the physiology in *D. melanogaster* with monoterpenes-treated *D. suzukii* flies.

## Material and methods

### Fly stocks

*Drosophila suzukii* was kindly provided by the Entomological Laboratory of the Agricultural Sciences Department of the University of Padua, (Italy) and maintained on an artificial diet with a 16:8 photoperiod, at a temperature of 22 ± 1 °C. *Drosophila melanogaster* mutant lines were as follows: TAR1^PL00408^ was generated by the Gene Disruption Project (Bloomington Stock Center, Indiana, USA) and TAR1-Gal4 was previously created in the Molecular Physiology group from the University of Kiel (El-Kholy et al., 2015). For behaviour experiments, *D. melanogaster y*^*1*^*w*^*1118*^ was used as a control. All *D. melanogaster* flies were raised on standard food at 25 ± 1 °C (12:12 light-dark photoperiod) as described previously (Li et al., 2016).

### Fumigant toxicity assay

A glass cylinder (10 cm in height, 4.5 cm inner diameter; 150 ml) was employed to calculate the monoterpenes LC_50_ values on *D. suzukii* and *D. melanogaster y*^*1*^*w*^*1118*^ and to perform the monoterpenes exposure. Monoterpenes including thymol, carvacrol, and α-terpineol were dissolved in acetone and applied to a filter paper (2 cm × 2 cm). The filter paper was placed on the bottom lid of the cylinder, inside a small cage to prevent direct contact of the flies with the monoterpenes. The concentrations ranged between 0.067 – 67 µl/L and acetone alone was used as negative control. After CO_2_ anesthetization, thirty flies (fifteen males and fifteen females) were placed inside the cylinder with 1 ml of solid diet. The top and the bottom of the cylinder were sealed with parafilm and the assay was maintained at 22 ± 1 °C for *D. suzukii* or 25 ± 1 °C for *D. melanogaster* flies. After 24 h the flies were collected. For the LC_50_ values calculation, at least one hundred flies were tested, in four replicates.

### Quantitative real-time PCR analysis

Total RNA was extracted from *D. suzukii* or *D. melanogaster y*^*1*^*w*^*1118*^ adult flies subjected to the monoterpene exposures using Aurum Total RNA Mini Kit (Bio-Rad, USA). One µg of RNA was treated with DNase I (Thermo Fisher, USA) and used for cDNA synthesis, carried out with the OneScript ® cDNA Synthesis Kit (Abm, Canada), according to the manufacturer’s instructions. Real time PCR was performed using a CFX Connect Real-Time PCR Detection System (Bio-Rad, USA) in a 12 µl reaction mixture containing 1.6 µl cDNA (diluted 1:2), 6 µl Sybr PCR Master Mix (Vazyme, China), 0.4 µl forward primer (10 µM), 0.4 µl reverse primer (10 µM) and 3.6 µl nuclease free water. Thermal cycling conditions were: 95 °C for 2 mins, 40 cycles at 95 °C for 15 s and 60 °C for 20 s. After the cycling protocol, a melting-curve analysis from 55 °C to 95 °C was applied. In *D. suzukii* expression of *TAR1* was normalized using *AK* and *TBP* genes that served as reference genes (Zhai et al., 2014). In *D. melanogaster y*^*1*^*w*^*1118*^ expression of *TAR1* was normalized using *actin* and *tubulin* genes that served as reference genes (Ponton et al., 2011). Gene-specific primers (**Table 1**) were used and four independent biological replicates, made in triplicate, were performed for each sample.

**Table 1.**
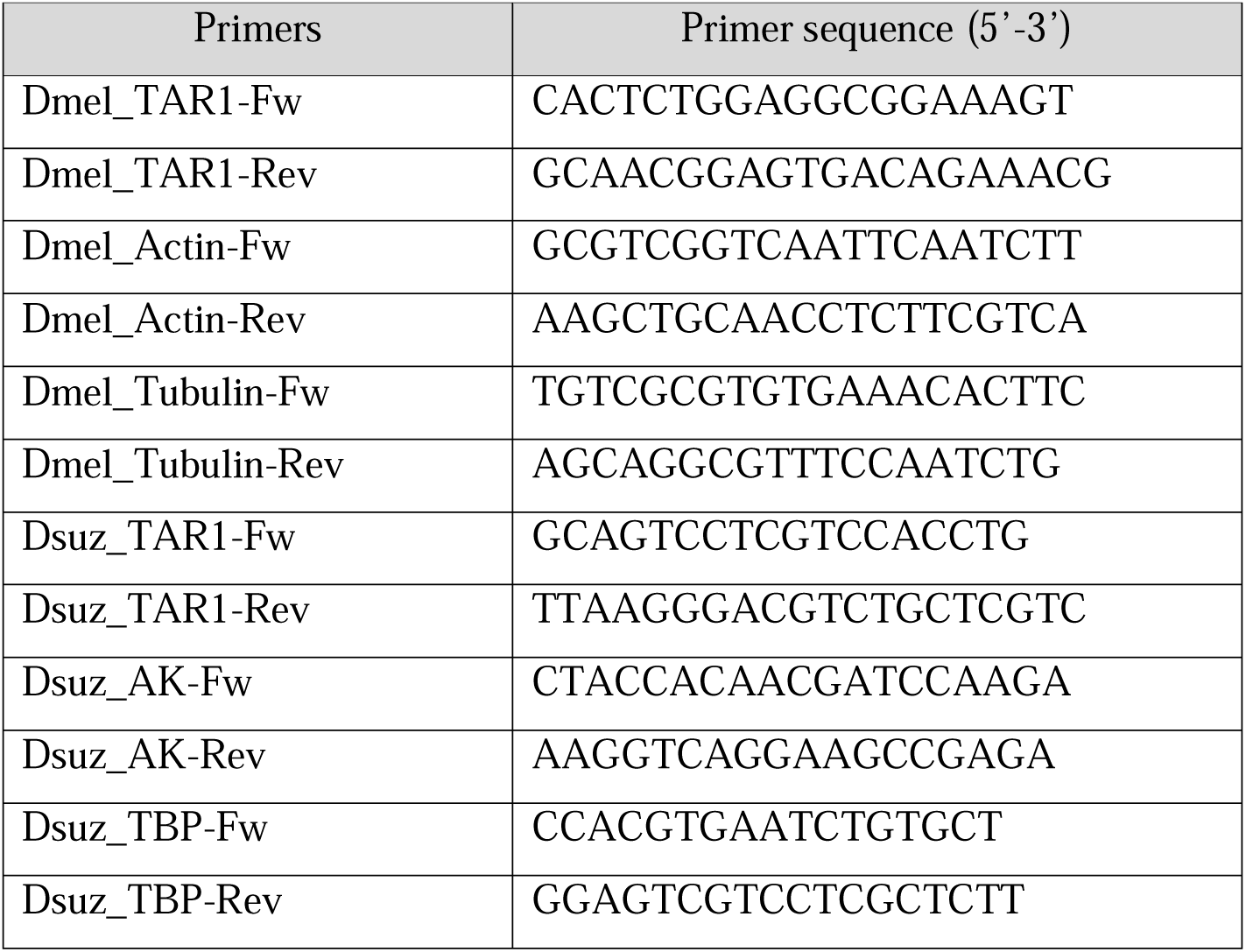
Primers used in this study.

### TAR1 immunohistochemistry

The TAR1-Gal4 *Drosophila* line was crossed with an UAS-GFP line in order to visualize the complete brain expression pattern of the receptor. The brains were dissected from F1 flies in cold Schneider’s *Drosophila* Medium and fixed in 4 % (w/v) paraformaldehyde in PBS for 90 mins at room temperature. The samples were then washed three times in PBST and blocked for 30 min in blocking buffer (1X PBS + 2 % NP-40 + 10 % goat serum) at room temperature. The samples were incubated with the primary antibodies in blocking buffer (anti-GFP rabbit 1:300 and anti-Nc82 mouse 1:20) overnight at 4 °C and washed three times for 5 min in PBST. Subsequently, the samples were incubated with the secondary antibodies in blocking buffer (donkey anti-rabbit IgG Alexa Fluor-488 1:300 and goat anti-mouse IgG Alexa Fluor 555 1:300) for 3 h at room temperature and washed twice for 5 min in PBST. Brains were mounted directly on slides and analysed by a Zeiss Axio Imager Z1 microscope equipped with an apotome (Zeiss, Germany).

### Body fat quantification

Total body triglyceride (TG) content was estimated using the Triglyceride (TG) colorimetric assay kit GPO-PAP method (Elabscience, China). Three flies were accurately weighted and homogenation medium (9 times the volume, phosphate buffer 0.1 mol/L, pH 7.4) was added. The sample was mechanically homogenized on ice with a motorized pestle and centrifugated (at 2500 rpm for 10 min). 7 µl of the supernatant were added to 700 µl of working solution kit, thoroughly mixed and incubated for 10 min at 37 °C in the dark. Absorbance was read at 510 nm and distilled water, added to 700 µl of working solution, was used as blank. Triglyceride content was estimated using a glycerol solution (2.26 mmol/L) as standard. Five independent biological replicates was performed for each sex and genotype.

### Dye-labelling food intake quantification

The dye-labelling food intake quantification was performed as described by Deshpande and co-workers (Deshpande et al., 2014), with minor modifications. In brief, five flies of each sex and genotype were placed into a vial with 2 ml of 1 X dyed medium (2.5 % yeast, 2.5 % sucrose, 1 % agar and 1 % Brilliant Blue FCF – Sigma Aldrich, USA). After 2 h of feeding, the flies were collected and frozen at −80 °C. Frozen flies were transferred to 1.5 ml Eppendorf tubes, homogenized with a manual pestle in 50 ul of 1 % PBST and centrifugated for 1 min at 12000 g to clear the debris. The supernatant absorbance was measured at 630 nm on a label-free EnSight Multimode Plate Reader (Perkin Elmer, USA). The values obtained from flies fed with non-labelled food were used as control and subtracted from experimental readings. To determine the dye concentration of each fly homogenate a standard curve was generated with serial dilutions of an initial 10 µl aliquot of the non-solidified dye-labelled food added to 990 µl of 1 % PBST. At least five independent biological replicates were performed for each sex and genotype.

### Metabolic rate determination

The measurement of the metabolic rate was assessed as described (Yatsenko et al., 2014). In brief, three adult flies were placed in each vial and the metabolic rate was measured for 2 h using the respirometry. The CO_2_ yield during the test was calculated based on the µl produced per h per fly. Data were obtained from five independent biological replicates.

### Rapid iterative negative geotaxis (RING) assay

The negative geotaxis assay was performed based on a published protocol (Gargano et al., 2005). In brief, five flies of each sex and genotype were placed into a 20 cm-tall glass tube without CO_2_-anaesthesia. The tube was tapped two times to move flies to the bottom and the climbing height of flies was photographed after 2 s. The average distance climbed in cm for each fly was measured using Image J software. Five independent biological replicates per sex and genotype were performed.

### Starvation resistance assay

The starvation resistance assay was performed placing twenty-five flies of each sex and genotype in vials containing 1% of agar. The vials were maintained at 22 ± 1 °C for *D. suzukii* or 25 ± 1 °C for *D. melanogaster*. Dead flies were counted every 2 h until all flies were dead. For each genotype and sex, four independent biological replicates were performed (at least one hundred flies).

### Statistical analyses

LC_50_ values were evaluated using POLO-plus software. All statistical analyses were performed using GraphPad Prism software (version 6). All data represent the mean values ± SEM, evaluated using the one-way ANOVA followed by Dunnett’s test for multiple comparisons.

## Results

### Monoterpenes LC_50_ calculation

The results of the LC50 estimation as obtained by POLO-plus analyses for each monoterpene, performed on both *D. suzukii* and *D. melanogaster y*^*1*^*w*^*1118*^ flies, are summarized in **Table 2**. The table reports the LC_50_-_90_ values, the 95% confidence limits (Robertson et al., 2017), the slopes (angular coefficients) of lines and the values of χ^2^ for each monoterpene.

**Table 2.** LC_50-90_ of fumigant active monoterpenes thymol, carvacrol and α-terpineol against *D. suzukii* and *D. melanogaster y*^*1*^ *w*^*1118*^.

| *D. suzukii* |  |  |  |  |
| --- | --- | --- | --- | --- |
| Compound | Slope (± SE) | LC<sub>50</sub> (95% CI) μl/L | LC<sub>90</sub> (95% CI) μl/L | $\chi^2$ |
| Thymol | 1.704 ± 0.318 | **1.085** (0.549 - 1.575) | 6.117 (4.362 – 10.854) | 2.605 |
| Carvacrol | 2.289 ± 0.341 | **0.844** (0.322 - 1.340) | 3.075 (1.930 – 8.744) | 3.991 |
| α-terpineol | 2.647 ± 0.307 | **1.494** (0.677 - 2.446) | 4.563 (2.754 – 14.164) | 6.493 |
| *D. melanogaster y<sup>l</sup> w<sup>1118</sup>* |  |  |  |  |
| Compound | Slope (± SE) | LC<sub>50</sub> (95% CI) μl/L | LC<sub>90</sub> (95% CI) μl/L | $\chi^2$ |
| Thymol | 1.749 ± 0.209 | **0.604** (0.152 – 2.036) | 3.260 (1.172 – 24.484) | 3.472 |
| Carvacrol | 1.864 ± 0.258 | **0.592** (0.156 – 1.636) | 2.888 (1.136 – 38.072) | 2.168 |
| α-terpineol | 1.677 ± 0.433 | **0.984** (0.300 – 1.524) | 5.252 (3.080 – 16.900) | 1.343 |

### TAR1 expression analysis after monoterpenes exposure

To evaluate the effect of the exposure to monoterpenes on the expression levels of *TAR1* gene in both *D. suzukii* and *D. melanogaster y*^*1*^*w*^*1118*^, flies were exposed to the LC_50_ concentrations of thymol, carvacrol and α-terpineol, respectively, and the mRNA levels analyzed by qPCR. The exposure induced an interesting downregulation of *TAR1* gene expression in both genotypes. In *D. suzukii*, significant differences were observed for thymol and carvacrol (**Figure 1, panel A**) but not for α-terpineol. On the other hand, in *D. melanogaster y*^*1*^*w*^*1118*^ all three monoterpenes induced a significant downregulation of *TAR1* although less marked as compared to *D. suzukii* (**Figure 1, panel B**).

**Figure 1.**
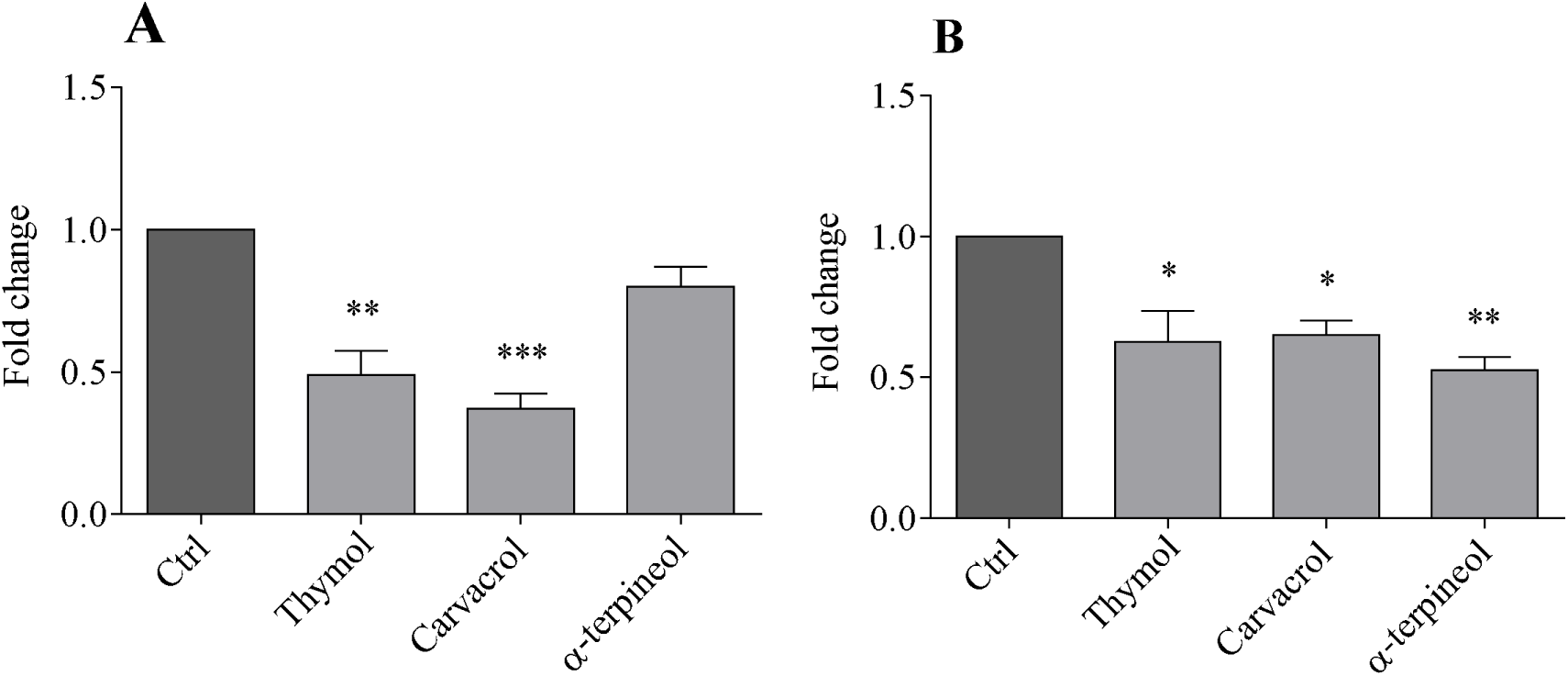
*D. suzukii* (panel A) and *D. melanogaster y*^*1*^*w*^*1118*^ (panel B) *TAR1* expression levels after 24 h of continuous exposure to the LC_50_ of thymol, carvacrol and α-terpineol. Data represent means ± SEM of four independent experiments performed in triplicate. *p < .05 **p < .01 ***p< .005 vs control according to one-way ANOVA followed by Dunnett’s test for multiple comparisons. Arginine kinase (*AK*) and TATA Box Protein (*TBP*) were used as reference genes in *D. suzukii* analysis (Zhai et al., 2014); *actin* and *tubulin* were used as reference gene in *D. melanogaster y*^*1*^*w*^*1118*^ analysis (Ponton et al., 2011).

### TAR1 expression in *D. melanogaster* brain

In order to determine the physiological functions controlled by TAR1, the receptor accumulation in *D. melanogaster* brains was investigated by immunohistochemistry. The Gal4-UAS system was used to selectively tag TAR1 with the GFP reporter protein, then recognized by the anti-GFP antibody. The receptor showed specific expression in the *pars intercerebralis* as well as lateral horn, sub-esophageal ganglia, mushroom bodies, and antennae mechanosensory -motor center (**Figure 2, panels A, B and C**), suggesting that TAR1 might be implicated in important physiological traits in *Drosophila*.

**Figure 2.**
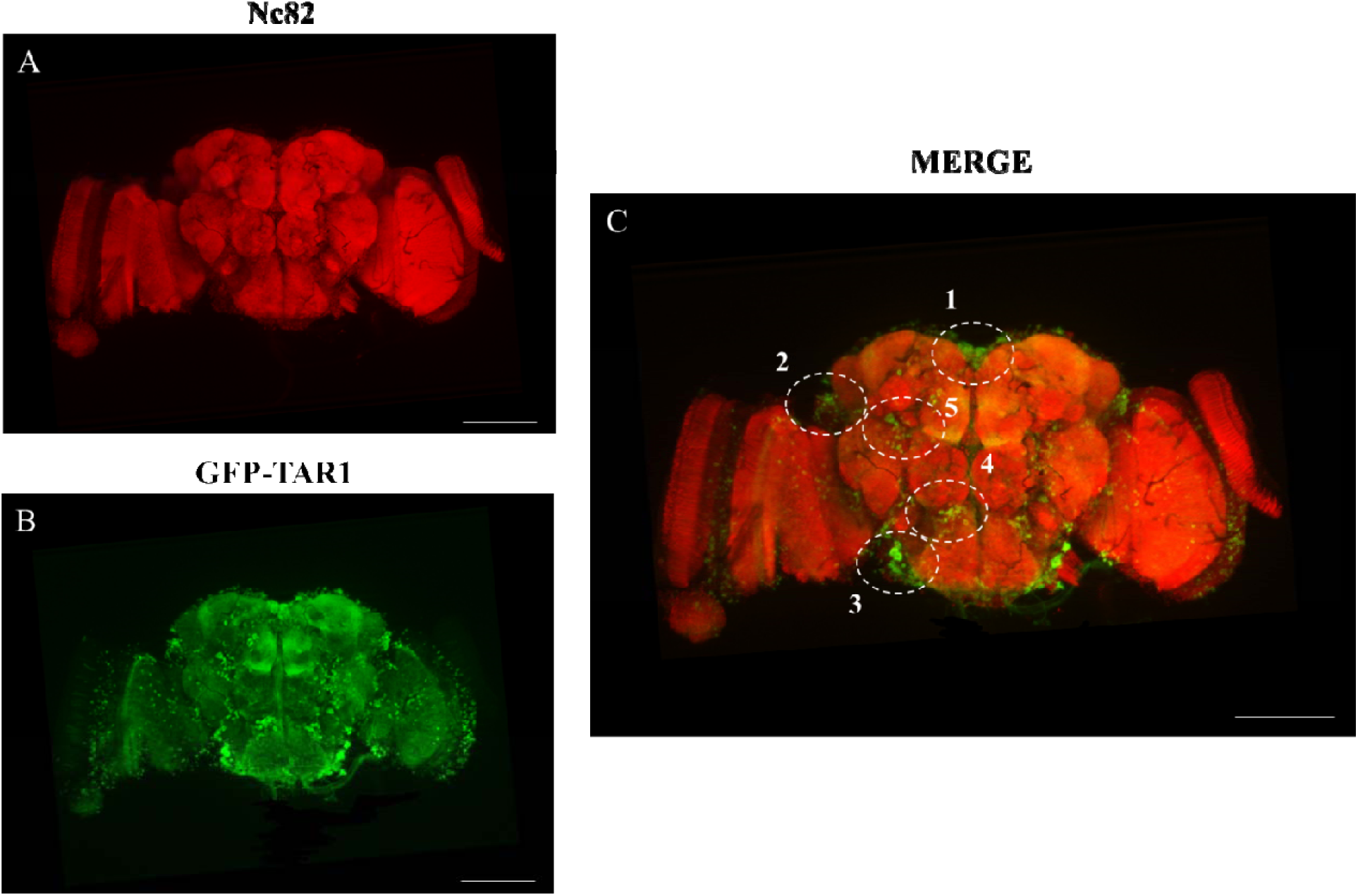
Activity of the TAR1 promoter in the *D. melanogaster* brain. Representative confocal images of GFP driven by TAR1-Gal4: synaptic regions are labelled with the presynaptic marker Nc82 (anti-Bruchpilot), TAR1 is marked by anti-GFP antibody. TAR1 is mainly localized in the *pars intercerebralis* (1), lateral horn (2), sub-esophageal zone (3), antennae mechanosensory – motor center (4) and mushroom bodies (5), as showed in the merge (**Panel C**). Scale bars = 100 µm for **A, B, C**.

### Role of TAR1 in *Drosophila* physiology

To elucidate the role of TAR1 in metabolic traits as well as locomotor control and physiological aspects in *Drosophila*, flies impaired in TAR1 (TAR1^PL00408^ or TAR1^−/−^) were enrolled in several behavioural assays. Flies with the same genetic background (*y*^*1*^*w*^*1118*^) were used as controls. In general, the absence of TAR1 translates into a higher propensity to triglycerides accumulation and food intake (**Figure 3, panels A and B**). Therefore, TAR1^−/−^ flies show higher resistance to starvation than control (**Figure 3, panel E**). These changes are furthermore associated with a slower metabolism in TAR1 impaired insects (**Figure 3, panel C**). The increased triglycerides accumulation and the slower metabolism could also be related to the lower propensity to movement of the TAR1^−/−^ flies (**Figure 3, panel D**).

**Figure 3.**
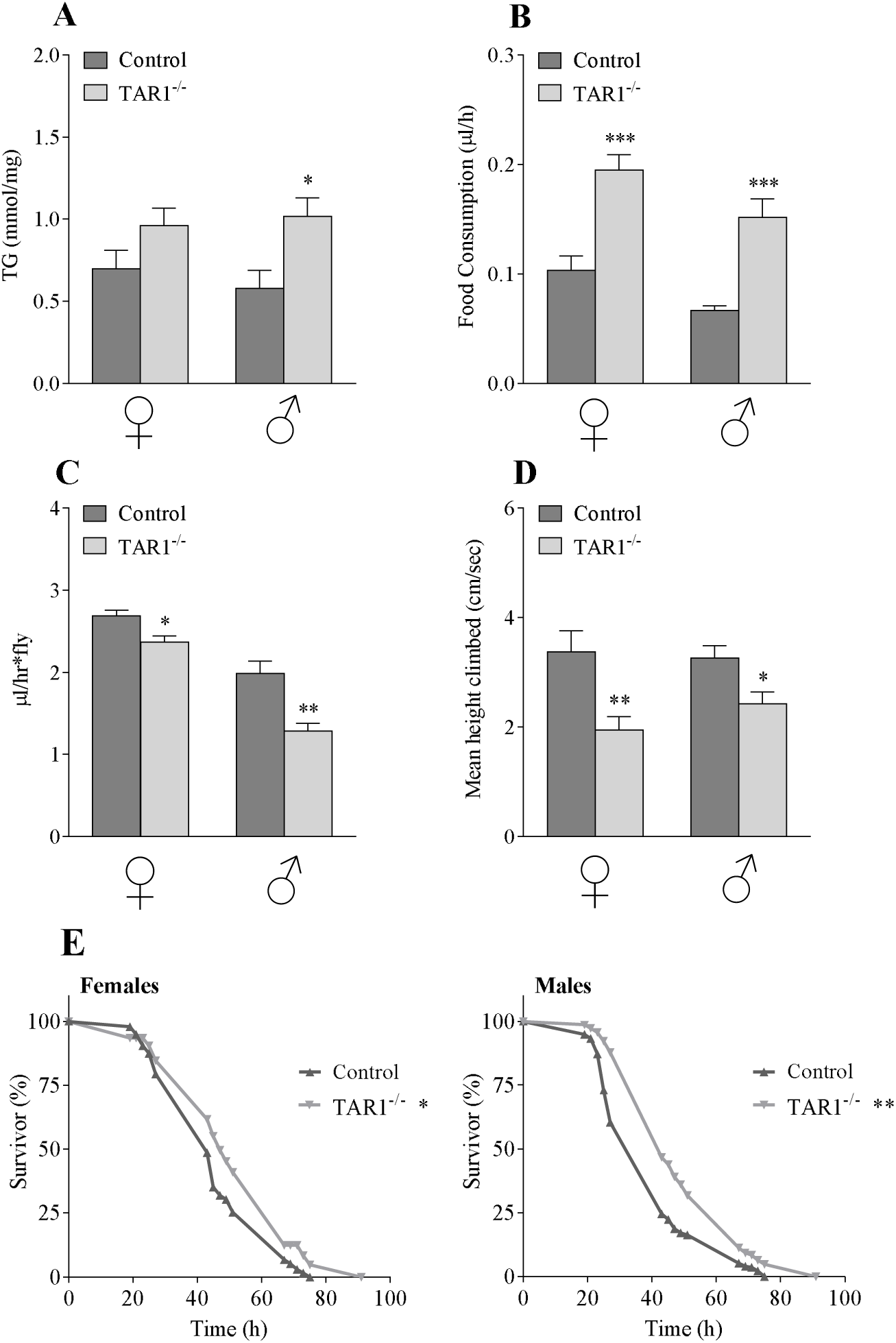
Physiological, metabolic and behavioural alterations in flies with an impaired TAR1. Total body triglyceride (TG) content (**panel A**), food intake quantification (**panel B**), metabolic rate (**panel C**), climbing activity measured by RING assay (**panel D**) and starvation resistance (**panel E**) were tested in control and TAR1^−/−^ animals of both sexes. For all experiments, means of at least four independent biological replicates ± SEM are shown. *p < .05 **p < .01 ***p< .005 vs control according to student’s *t*-test. In starvation resistance, statistical analyses were performed using the log-rank test.

To test whether monoterpenes, besides downregulationg *TAR1*, might also alter the physiology of *D. suzukii* and *D. melanogaster* (wild type or TAR1-/-), flies 24 h after the continued monoterpenes LC_50_ exposure were challenged with several behavioural tests.

### Monoterpenes treatment – effects on total body triglyceride (TG) content

24 h of exposure to monoterpenes caused a higher TG content in males of both *D. suzukii* and *D. melanogaster y*^*1*^*w*^*1118*^ flies as compared to females (**Figure 4**). In particular, the TG content was significantly higher upon thymol and carvacrol exposure, only in *D. suzukii* males (**Figure 4, panel B**), while, both *D. melanogaster y*^*1*^*w*^*1118*^ females and males showed a significantly higher TG content after carvacrol exposure (**Figure 4, panels C and D**). When the same treatments were applied to *D. melanogaster* TAR1^−/−^ insects, no changes were observed in TG content, which was indistinguishable from the untreated control sample. This evidence would suggest that monoterpenes can induce an increase in total fat deposition that requires TAR1 receptors be functional (**Figure 4, panels E and F**).

**Figure 4.**
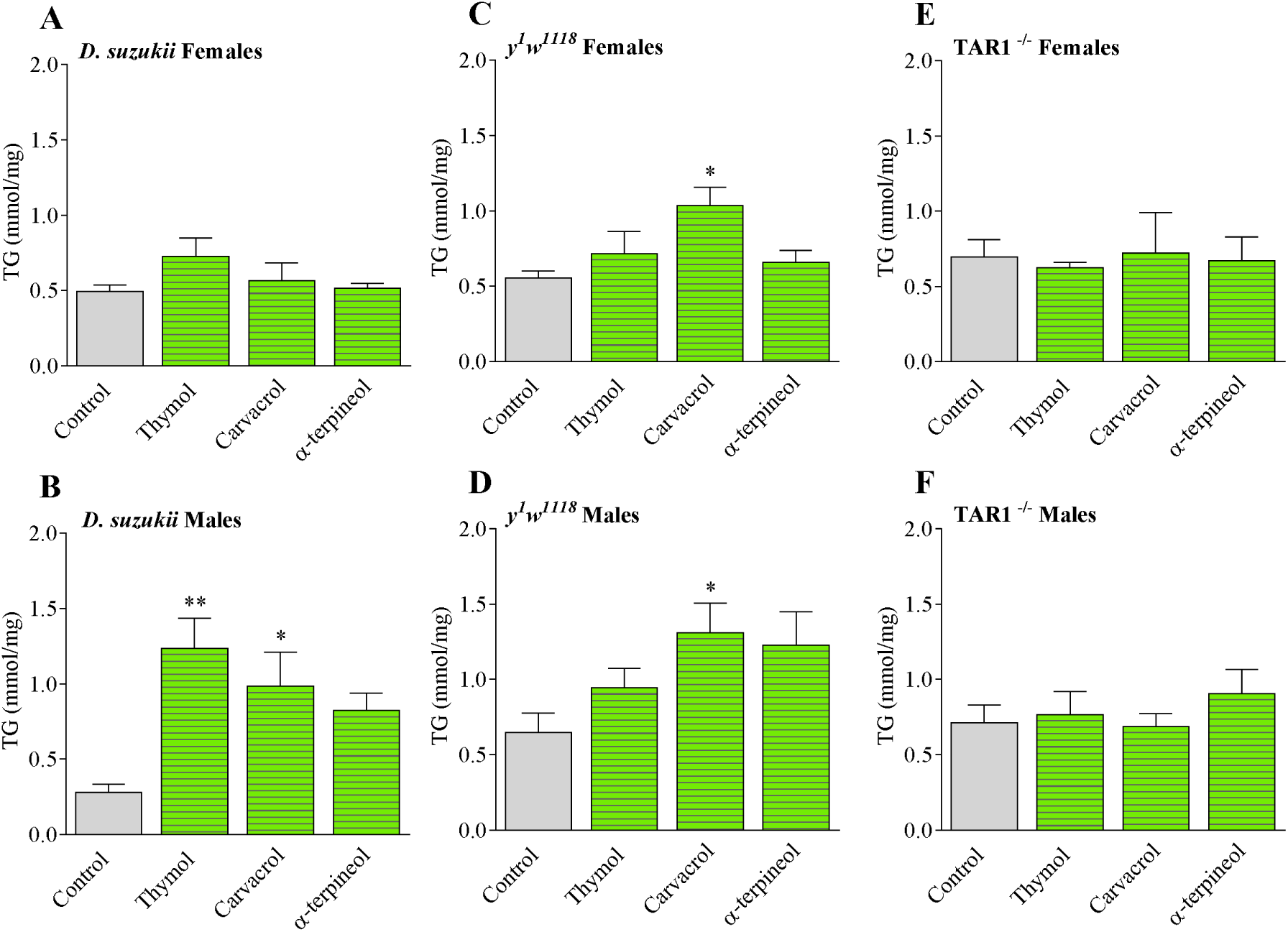
Total body triglyceride (TG) content after 24 h of exposure to monoterpenes, in *D. suzukii* (panels A and B), *D. melanogaster y*^*1*^*w*^*1118*^ (panels C and D) and *D. melanogaster* TAR1^−/−^ (panels E and F). Data shown are the means ± SEM of four independent biological replicates. *p < .05 **p < .01 vs control according to one-way ANOVA followed by Dunnett’s test for multiple comparisons.

### Monoterpenes treatment – effects on food intake

The food consumption was quantified after two hours of feeding on a dye-labelled diet. A significantly high food intake was observed only after α-terpineol exposure in both *D. suzukii* and *D. melanogaster y*^*1*^*w*^*1118*^ of both sexes (**Figure 5, panels A, B, C and D**). The increased food intake might explain the high triglyceride levels observed in both *D. suzukii* and *D. melanogaster y*^*1*^*w*^*1118*^ sexes after monoterpenes exposure. On the other hand, the monoterpene treatments did not cause any change in food consumption in *D. melanogaster* TAR1^−/−^ mutant flies (**Figure 5, panels E and F**) further suggesting the requirement for an active TAR1.

**Figure 5.**
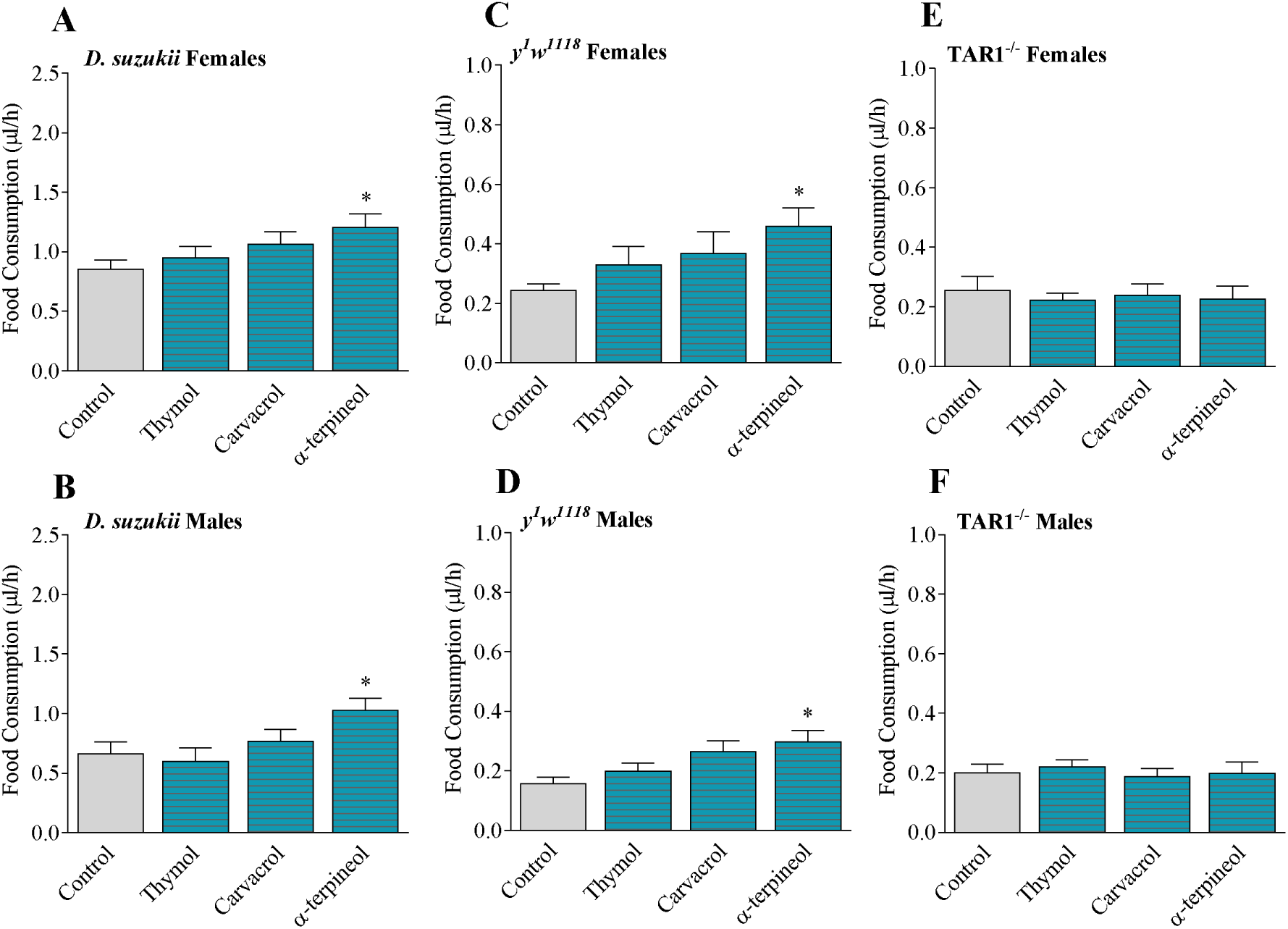
Food intake, after 24 h of exposure to monoterpenes, in *D. suzukii* (panels A and B), *D. melanogaster y*^*1*^*w*^*1118*^ (panels C and D) and *D. melanogaster* TAR1^−/−^ (panels E and F) measured as μl of diet per hour. Data shown are the means ± SEM of five independent biological replicates. *p < .05 vs control according to one-way ANOVA followed by Dunnett’s test for multiple comparisons.

### Monoterpenes treatment – effects on metabolic rate

In order to determine if the monoterpenes and the TAR1 downregulation might affect the metabolism, the metabolic rate was analysed in all *D. suzukii* and *D. melanogaster* genotypes after treatment with the different monoterpenes. In *D. suzukii*, only males treated with the three monoterpenes showed a significantly lower metabolic rate than control flies (**Figure 6, panels A and B**). Carvacrol and α-terpineol were able to reduce the metabolic rate in *D. melanogaster y*^*1*^*w*^*1118*^ males and females as well (**Figure 6, panels C and D**). Conversely, *D. melanogaster* TAR1^−/−^ metabolic rate appeared unaffected by the treatments therefore undistinguishable from that of the untreated controls (**Figure 6, panels E and F**).

**Figure 6.**
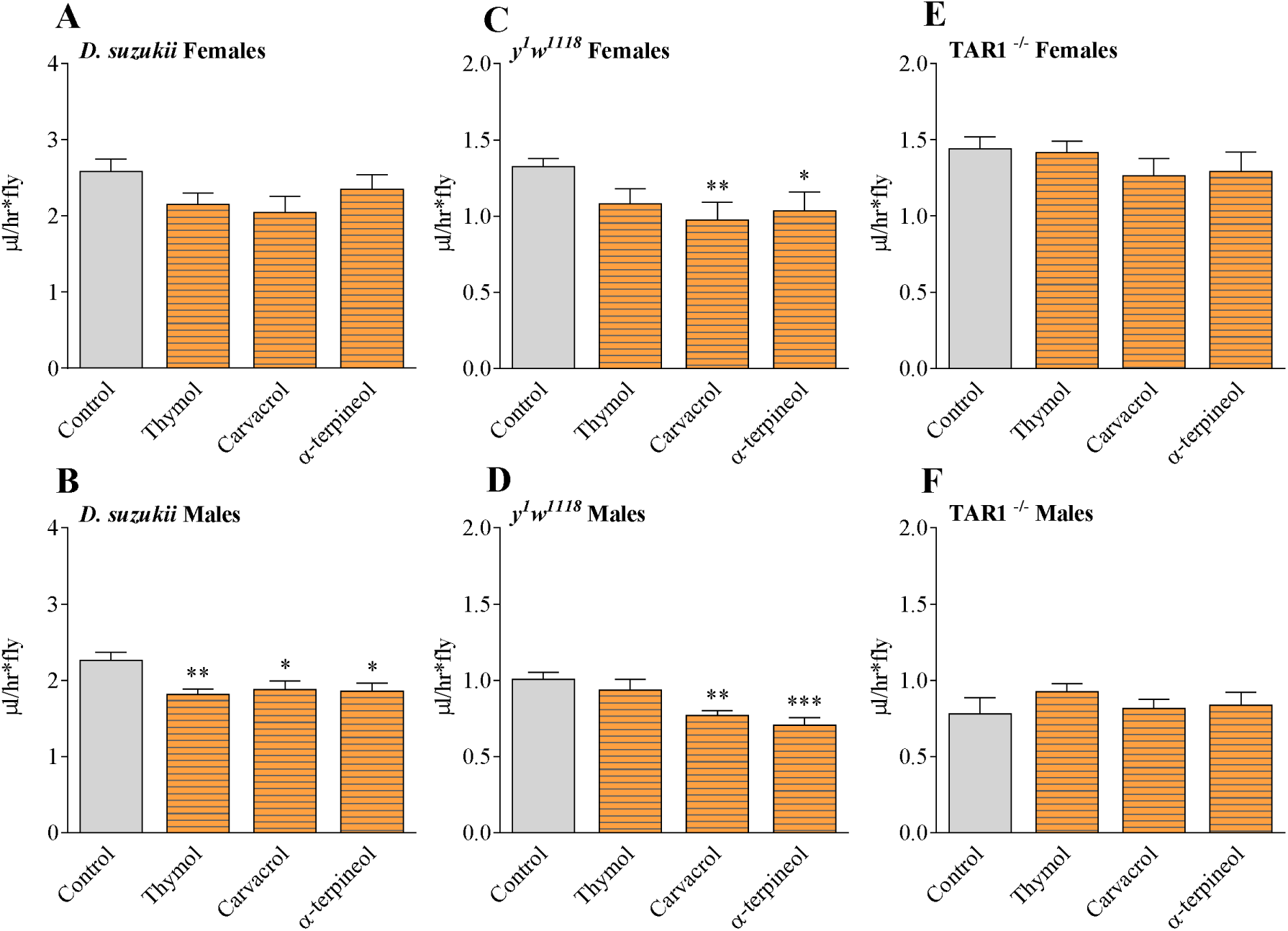
Metabolic rate, after 24 h of exposure to monoterpenes, in *D. suzukii* (panels A and B), *D. melanogaster y*^*1*^*w*^*1118*^ (panels C and D) and *D. melanogaster* TAR1^−/−^ (panels E and F). Data shown are the means ± SEM of five independent biological replicates. *p < .05 **p < .01 ***p< .005 vs control according to one-way ANOVA followed by Dunnett’s test for multiple comparisons.

### Monoterpene treatment – effects on locomotory activity

The observed metabolic changes in terms of energy expenditure and TG content might also affect flies physical activities. Therefore, the ability of flies exposed to monoterpenes to walk upwards on a vertical surface in negative geotaxis was used as a motility behavioural assay. In comparison to controls, *D. suzukii* and *D. melanogaster y*^*1*^*w*^*1118*^ males showed a statistically significant reduction in climbing ability only after α-terpineol treatment (**Figure 7, panels B and D**). *D. melanogaster y*^*1*^*w*^*1118*^ females motility was negatively affected only by thymol (**Figure 7, panel C**), while *D. suzukii* females did not respond to the RING assay at all, in both control and treated samples (**Figure 7, panel A**). The climbing ability in both *D. melanogaster* TAR1^−/−^ sexes was unaffected by the exposure to monoterpenes, confirming the hypothesis of TAR1 involvement in this behavioural trait.

**Figure 7.**
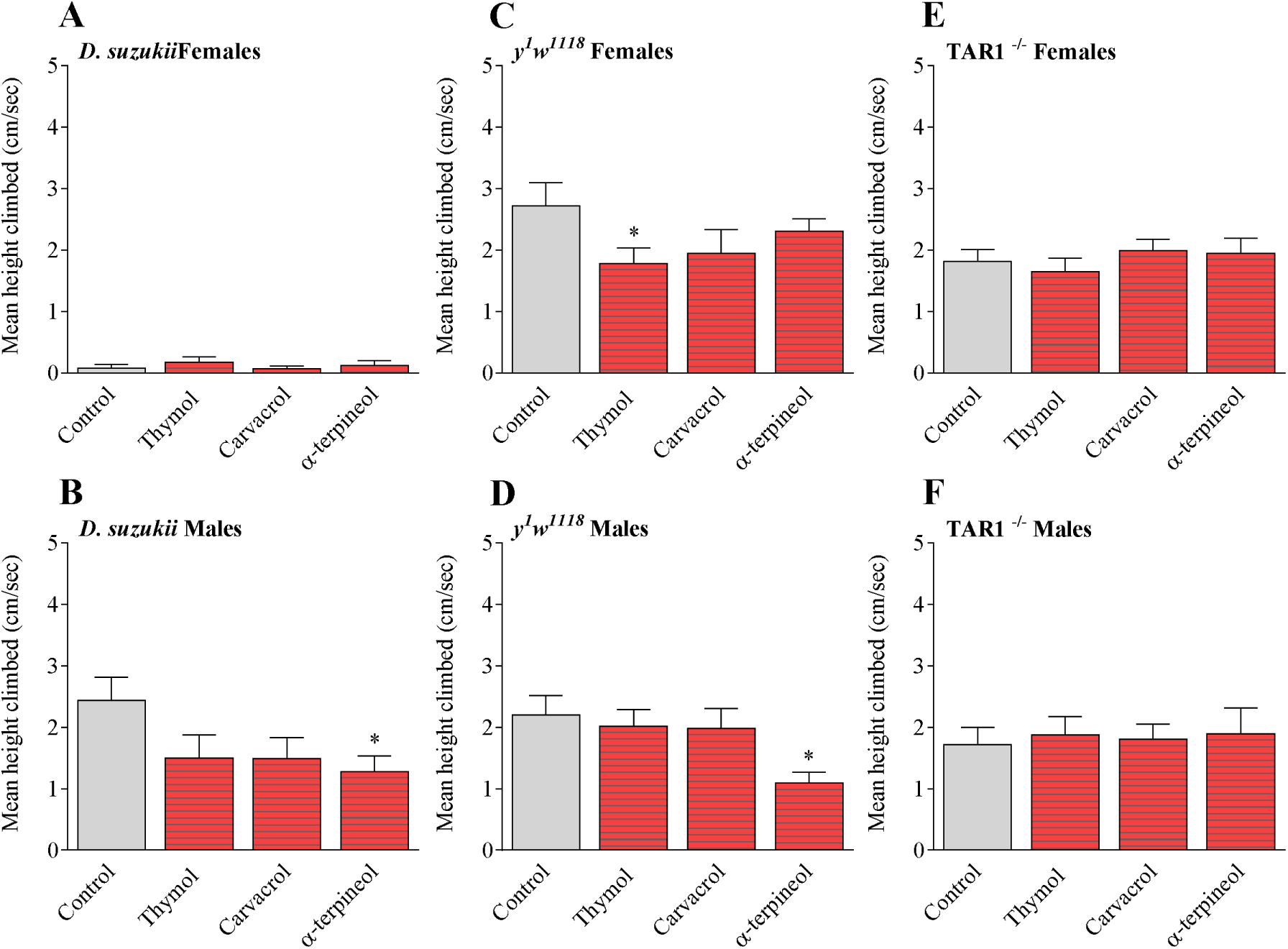
RING assay, after 24 h of exposure to monoterpenes, on *D. suzukii* (panels A and B), *D. melanogaster y*^*1*^*w*^*1118*^ (panels C and D) and *D. melanogaster* TAR1^−/−^ (panels E and F). The vertical movement capacity for each insect is expressed in cm per second. Data shown are the means ± SEM of five independent biological replicates. *p < .05 vs control according to one-way ANOVA followed by Dunnett’s test for multiple comparisons.

### Monoterpene treatment – effects on starvation resistance

Finally, a starvation resistance assay was performed to investigate whether the monoterpene-mediated metabolic modifications could affect the general fitness. Given the higher food intake and TG content caused by the treatment, an enhanced starvation resistance was expected. *D. suzukii* and *D. melanogaster y*^*1*^*w*^*1118*^ showed different results depending on the monoterpene used as compared to control (**Figure 8, panels A, B, C and D**). According to log-rank statistical analysis, a significant reduction in starvation resistance was detected in *D. suzukii*, both males and females, after carvacrol treatment (**Figure 8, panels A and B**) while both *D. melanogaster y*^*1*^*w*^*1118*^ sexes were less resistant to starvation after thymol exposure. Moreover, α-terpineol treatment reduced starvation resistance only in *D. melanogaster y*^*1*^*w*^*1118*^ females flies (**Figure 8, panels C and D**). Conversely, the carvacrol exposure significantly increased the starvation resistance in *D. melanogaster y*^*1*^*w*^*1118*^ males (**Figure 8, panel C**). *D. melanogaster* TAR1^−/−^ mutant were again unaffected by the treatment, thus showing starvation resistance comparable to controls (**Figure 8, panels E and F**).

**Figure 8.**
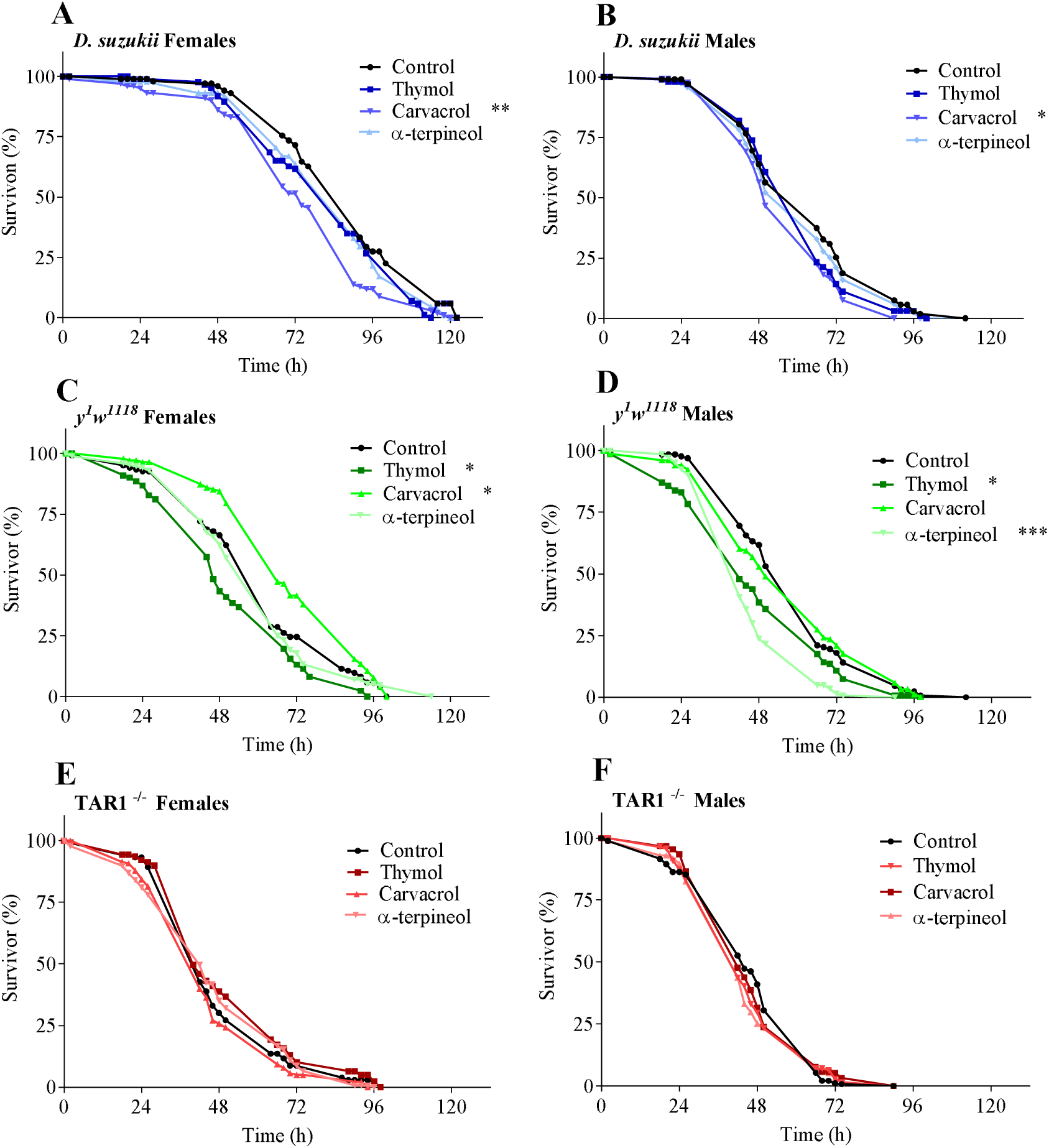
Starvation resistance, after 24 h of exposure to monoterpenes, on *D. suzukii* (panels A and B), *D. melanogaster y*^*1*^*w*^*1118*^ (panels C and D) and *D. melanogaster* TAR1^−/−^ (panels E and F). Five independent biological replicates were performed with the log-rank test statistical analysis. *p < .05, **p<.01, ***p<.005 vs control.

## Discussion

The biogenic amine TA is a mediator of several physiological functions in invertebrates (Roeder, 2005; Lange, 2009), but its mechanism of action is still far from being fully characterized. TA activates intracellular responses by interacting with specific GPCRs, the tyramine receptors TAR (Saudou et al., 1990; Roeder et al., 2003). TAR1 is highly expressed in the CNS of numerous insects, thus suggesting its involvement in essential behavioural processes (El-Kholy et al., 2015; Hana & Lange, 2017; Finetti et al., 2020). Furthermore, several studies showed that TAR1 could be a direct target for biomolecules with insecticidal action, such as monoterpenes. In fact, it has been reported that the *D. melanogaster* and *R. microplus* TAR1s, when expressed in a heterologous cell system, respond to the administration of monoterpenes with an increased release of cytosolic calcium (Enan, 2005a; Gross et al., 2017). Recently, the same intracellular response has been observed in our laboratory for *D. suzukii* TAR1, allowing to hypothesize that the interaction between monoterpene and receptor causes a downregulation of the gene coding for the receptor (Finetti et al., 2020). To further study the effects of the monoterpenes on TAR1 and on the insect physiology, a *D. melanogaster* TAR1 deficient line (TAR1^−/−^) was evaluated together with matching controls and *D. suzukii*. Comparative studies using these two *Drosophila* species are possible since they are phylogenetically highly related and their TAR1 share a high degree of homology (98 %) (Finetti et al., 2020).

Firstly, the identification of the LC_50_ for the three monoterpenes thymol, carvacrol and α-terpineol, for both *D. suzukii* and *D. melanogaster y*^*1*^*w*^*1118*^ via a fumigant assay (Park et al., 2016), revealed that the most toxic monoterpene was carvacrol with a LC_50_ of 0.844 µl/L for *D. suzukii* and 0.592 µl/L for *D. melanogaster*. Similarly, Zhang and co-workers (2016) observed that carvacrol was the most toxic monoterpene for *D. melanogaste*r. Interestingly, when TAR1^−/−^ flies were treated with the monoterpenes at the LC_50_ calculated for the *y*^*1*^*w*^*1118*^ strain a 40 % reduced mortality was observed as compared to the control (data not shown), suggesting a strong correlation between TAR1 and the insecticidal activity of these monoterpenes. A similar observation was made in a *D. melanogaster* TAR1 deficient strain (specifically TyrR^Neo30^), which appeared to be insensitive to thymol and carvacrol when topically applied (Enan, 2005a).

All three monoterpenes tested, thymol, carvacrol and α-terpineol, after 24 h of fumigant treatment, were able to induce a TAR1 downregulation not only in *D. suzukii* (as already established, Finetti et al., 2020) but also in *D. melanogaster*. Since TAR1 is mainly expressed in the CNS, the greatest impact of its downregulation might be expected in this region.

As shown by El-Kholy et al. (2015), in a study focused on *D. melanogaster* brain, TAR1 is expressed in the *pars intercerebralis*, mushroom bodies and ellipsoid body, as confirmed also by Li et al. (2016). Our study revealed that TAR1 is strongly expressed not only in the *pars intercerebralis* and the mushroom bodies but also in lateral horn, sub-esophageal ganglia, and antennae mechanosensory centre. Even if the physiological significance of these specific TAR1 expression patterns in the *Drosophila* SNC is still unclear, they are likely directly connected to the functions associated with the corresponding brain areas. The *pars intercerebralis* is an important insect neuroendocrine center composed by neurosecretory cells that regulate feeding (olfactory/gustatory perception of food sources; feedback information from the intestinal tract and body cavity regarding the urgency of feeding) and reproductive behaviours (Velasco et al., 2006). TAR1^−/−^ mutant flies showed a phenotypic profile that correlates with these observations. These flies are in fact characterized by increased body fat, higher food intake and starvation resistance as well as reduced locomotor activity and metabolic rate in comparison to *y*^*1*^*w*^*1118*^ controls (Li et al., 2016; Li et al., 2017). These metabolic alterations were not sex dependent, although the effects in TAR1^−/−^ males appeared to be more pronounced as compared to those seen in females. This could be related to sex-dependent differences in TAR1 expression, whose mRNAs accumulated at higher levels in males than in females (Finetti et al., 2020). Despite all this, little is still known on the precise mechanism by which the tyraminergic system modulates essential metabolic traits such as fat body, food intake, starvation resistance, locomotor activity and metabolic rate.

In insects, fat is mainly stored in the fat body, which is, at the same time, one of the most important metabolic centers (Arrese & Soulages, 2010). Lipid storage and release are mainly controlled by two hormones, the *Drosophila* insulin-like peptides (mainly dILP2) and the AKH (Adipokinetic hormone, analogous to the mammalian glucagon) (Roeder, 2020). During an acute stress situation, the mobilization of lipids is essential for survival. This mechanism appears to be also controlled by both, OA and TA, presumably through modulation of dILP secretion (Fields & Woodring, 1991; Orchard et al., 1993). In fact, it has recently been observed that in *C. elegans*, during acute stress, TA accumulates, which in turn modulates insulin signal (De Rosa et al., 2019). Therefore, increased TG level observed in TAR1^−/−^, as compared to *y*^*1*^*w*^*1118*^ control flies, might be related to a direct tyraminergic action on the release of dILPs. RNAi-mediated TAR1 silencing, targeted to the fat body, triggered reduction of dILP2 in insulin-producing cells in the *D. melanogaster pars intercerebralis* and an increased TG accumulation (Li et al., 2017). The increased TG levels in TAR1^−/−^ flies could also be linked to enhanced food intake as well as to lower movement propensity and metabolic rate. It has recently been proposed, in fact, that TAR1 could be involved in processes related to sugar sensibility and food intake regulation (Ishida & Ozaki, 2010). For example, both *honoka* and TAR1 KO flies (TyR^f05682^) showed a reduced sugar response (Damrau et al., 2019) linked to differences in food intake. It is worth noting that TAR1 is highly expressed in neurons located in the sub-esophageal ganglia that are presumably associated with the salivary glands and neck muscles control, thus linked with feeding.

After monoterpene treatments, both *D. melanogaster y*^*1*^*w*^*1118*^ and *D. suzukii* showed alterations in all behavioural assays performed. The link between monoterpene treatment and TAR1 downregulation is supported by the higher food intake observed in response to this treatment. When the *D. melanogaster* TAR1^−/−^ deficient line was considered, no phenotypic changes were observed whatsoever after exposure to monoterpenes, suggesting that the alterations observed in the other genotypes require the correct expression of a functioning receptor. This further confirms the relationship between monoterpenes-induced behavioural changes and TAR1. TAR1-mediated physiological alterations due to monoterpenes were also observed in *P. regina*. In fact, D-limonene treatment decreased TA levels in *P. regina* brain, causing a direct modification of the food intake (Nishimura et al., 2005). This different response to food stimuli was subsequently attributed to a probable alteration of the TAR1 expression at the level of the sub-exophageal ganglion (Yshida & Ozaki, 2011). Furthermore, thymol and carvacrol appeared to play a crucial role modulating ant behaviour (locomotion and aggression), through aminergic regulation (Mannino et al., 2018).

In conclusion, this study shows that monoterpenes might be instrumental in the manipulation of the insect behaviour via TAR1. In fact, sublethal concentrations of thymol, carvacrol and α-terpineol downregulate TAR1 expression, ultimately affecting important metabolic traits such as starvation resistance and energy storage. Moreover, this work demonstrated that monoterpenes, in addition to their insecticidal properties, can modify the metabolism and fitness of surviving *D. suzukii* opening to innovative applications of these molecules in the pest control.

## Acknowledgements

We would like to thank Dr. Morena de Bastiani (University of Ferrara) for excellent technical assistance and Dr. Federica Albanese (University of Ferrara, Italy) for linguistic improvement of the manuscript.

## References

Arrese, E.L. and Soulages, J.L. (2010). Insect fat body: energy, metabolism, and regulation. Annual Review of Entomology 55, 207–225.

Asplen, M.K., Anfora, G., Biondi, A., Choi, D.S., Chu, D., Daane, K.M. and Desneux, N. (2015). Invasion biology of spotted wing Drosophila (*Drosophila suzukii*): a global perspective and future priorities. Journal of Pest Science 88(3), 469–494.

Audsley, N. and Dom, R.E. (2015). G protein coupled receptors as target for next generation pesticides. Insect Biochemistry and Molecular Biology 67, 27–37.

Bakkali, F., Averbeck, S., Averbeck, D. and Idaomar, M. (2008). Biological effects of essential oils – a review. Food and Chemical Toxicology 46, 446–475.

Blenau, W., Rademacher, E. and Baumann, A. (2011). Plant essential oils and formamidines as insecticides/acaricides: What are the molecular targets? Apidologie 43(3), 334–347.

Brembs, B., Christiansen, F., Pfluger, H.J. and Duch, C. (2007). Flight initiation and maintenance deficits in flies with genetically altered biogenic amines levels. Journal of Neuroscience 27, 11122–11131.

Brigaud, L., GrosmaÎtre, X., Franîois, M.C. and Jacqion-Joly, E. (2009). Cloning and expression pattern of a putative octopamine/tyramine receptor in antennae of the noctuid moth *Mamestra brassicae*. Cell Tissue Research 335, 445–463.

Cini, A., Ioriatti, C. and Anfora, G. (2012). A review of the invasion of *Drosophila suzukii* in Europe and a draft research agenda for integrated pest management. Bulletin of Insectology 65(1), 149–160.

Dam, D., Molitor, D. and Beyer, M. (2019). Natural compounds for controlling *Drosophila suzukii*. A review. Agronomy for Sustainable Development 39, 53.

Damrau, C., Toshima, N., Tanimura, T., Brembs, B. and Colomb, J. (2019). Octopamine and tyramine contribute separately to the counter-regulatory response to sugar deficit in Drosophila. Frontiers in Systems Neuroscience 11, 100.

De Rosa, M.J., Veuthey, T., Florman, J., Grant, J., Blanco, M.G., Andersen, N., Donnelly, J., Rayes, D. and Alkema, M.J. (2019). The flight response impairs cytoprotective mechanism by activating the insulin pathway. Nature 573, 135–138.

Deshpande, S.A., Carvalho, G.B., Amador, A., Phillips, A.M., Hoxha, S., Lizotte, K.J. and Ja, W.W. (2014). Quantifying *Drosophila* food intake: comparative analysis of current methodology. Nature Methods 11(5), 535–540.

Desneux, A., Decourtye, A. and Delpuech, J.M. (2007). The sublethal effects of pesticides on beneficial arthropods. Annual Review of Entomology 52, 81–106.

Donini, A. and Lange, A.B. (2004). Evidence for a possible neurotransmitter/neuromodulator role of tyramine on the locust oviducts. Journal of Insect Physiology 50, 351–361.

Duportets, L., Barrozo, R., Bozzolan, F., Gaertner, C., Anton, S., Gadenne, C. and Debernard, S. (2010) Cloning of an octopamine/tyramine receptor and plasticity of its expression as a function of adult sexual maturation in the male moth *Agrotis ipsilon*. Insect Molecular Biology 19(4), 489–499.

El-Kholy, S., Stephano, F., Li, Y., Bhandari, A., Fink, C. and Roeder, T. (2015). Expression analysis of octopamine and tyramine receptors in *Drosophila*. Cell and Tissue Research 361(3), 669–684.

Enan, E.E. (2001). Insecticidal activity of essential oils: octopaminergic sites of action. Comparative Biochemistry and Physiology – Part C: Toxicology & Pharmacology 130, 325–337.

Enan, E.E. (2005a). Molecular response of *Drosophila melanogaster* tyramine receptor cascade to plant essential oils. Insect Biochemistry and Molecular Biology 35, 309–321.

Enan, E.E. (2005b). Molecular and pharmacological analysis of an octopamine receptor from American cockroach and fruit fly in response to plant essential oils. Archives of Insect Biochemistry and Physiology 59, 161–171.

Evans, P.D. and Maqueira, B. (2005). Insect octopamine receptors: a new classification scheme based on studies of cloned G-protein coupled receptors. Invertebrate Neuroscience 5, 111–118.

Fields, P.E. and Woodring, J.P. (1991). Octopamine mobilization of lipids and carbohydrates in the house cricket, *Acheta domesticus*. Journal of Insect Physiology 37(3), 193–199.

Finetti, L., Ferrari, F., Calò, G., Cassanelli, S., De Bastiani, M., Civolani, S. and Bernacchia G. (2020). Modulation of *Drosophila suzukii* type 1 tyramine receptor (DsTAR1) by monoterpenes: a potential new target for next generation biopesticides. Pesticide Biochemistry and Physiology 165, 91–101.

Fuchs, S., Behrends, V., Bundy., J.G., Crisanti, A. and Nolan, T. (2014). Phenylalanine metabolism regulates reproduction and parasite melanisation in the malaria mosquito. PLoS ONE 9, e84865.

Gargano, J.W., Martin, I., Bhandari, P. and Grotewiel, M.S. (2005). Rapid iterative negative geotaxis (RING): a new method for assessing age-related locomotor decline in *Drosophila*. Experimental Gerontology 40(5), 386–395.

Gross, A.D., Temeyer, K.B., Day, T.A., Pérez de León, A.A., Kimber, M.J. and Coats J.R. (2017). Interaction of plant essential oil terpenoids with the southern cattle tick tyramine receptor: A potential biopesticide target. Chemico-Biological Interactions 263, 1–6.

Hana, S. and Lange, A. (2017). Cloning and functional characterization of Octβ2-receptor and Tyr1-receptor in the Chagas disease vector, *Rhodnius prolixus*. Frontiers in Physiology 8, 744.

Hardie, S.L., Zhang, J.X. and Hirsh, J. (2007). Trace amines differentially regulate adult locomotor activity, cocaine sensibility and female fertility in *Drosophila melanogaster*. Developmental Neurobiology 67, 1396–1405.

Haviland, D.R. and Beers, E.H. (2012). Chemical control programs for *Drosophila suzukii* that comply with international limitations on pesticides residues for exported sweet cherries. Journal of Integrated pest Management 3(2), 1–6.

Hirashima, A., Yamaji, H., Yoshizawa, T., Kuwano, E. and Eto, M. (2007). Effect of tyramine and stress on sex-pheromone production in pre- and post-mating silkworm moth, *Bombyx mori*. Journal of Insect Physiology 53, 1242–1249.

Houghton, P.J., Ren, Y. and Howes, M.J. (2006). Acetylcholinesterase inhibitors from plants and fungi. Natural Product Reports 23(2), 181–199.

Ishida, Y. and Ozaki, M. (2011). A putative octopamine/tyramine receptor mediating appetite in a hungry fly. Naturwissenschaften 98, 635–638.

Isman, M.B. (2020). Botanical insecticides in the twenty-first century – fulfilling their promise? Annual Review of Entomology 65, 233–249.

Jankowska, M., Wyszkowska, J., Stankiewicz, M and Rogalska, J. (2018). Molecular targets for components of essential oils in the insect nervous system – a review. Molecules 23, 34.

Jensen, H.R., Scott, I.M., Sims, S.R., Trudeau, V.L. and Arnason, J.T. (2006). The effect of a synergistic concentration of a Piper nigrum extract used in conjunction with pyrethrum upon gene expression in *Drosophila melanogaster*. Insect Molecular Biology 15, 329–339.

Kim, J., Jang, M., Shin, E., Kim, J., Lee, S. H. and Park, C. G. (2016). Fumigant and contact toxicity of 22 wooden essential oils and their major components against *Drosophila suzukii* (Diptera: Drosophilidae). Pesticide Biochemistry and Physiology 133, 35–43.

Konstantopoulou, I., Vassipoulou, L., Mauragani-Tsipidov, P. and Scouras, Z.G. (1992). Insecticidal effects of essential oils. A study of the effects of essential oils extracted from eleven Greek aromatic plants on *Drosophila auraria*. Experientia 48, 616–619.

Kostyukovsky, M., Rafaeli, A., Gileadi, C., Demchenko, N. and Shaaya, E. (2002). Activation of octopaminergic receptors by essential oil constituents isolated from aromatic plants: possible mode of action against insect pests. Pest Management Science 58(11), 1101–1106.

Kutsukake, M., Komatsu, A., Yamamoto, D. and Ishiwa-Chigusa, S. (2000) A tyramine receptor gene mutation causes a defective olfactory behaviour in *Drosophila melanogaster*. Gene 245, 31–42.

Lange, A.B. (2009). Tyramine: from octopamine precursor to neuroactive chemical in insects. General and Comparative Endocrinology 162, 18–26.

Lee, J.C., Bruck, D.J., Curry, H., Edwards, D., Haviland, D.R., Van Steenwyk, R.A. and Yorgey, B.M. (2011). The susceptibility of small fruits and cherries to the spotted-wing Drosophila, *Drosophila suzukii*. Pest Management Science 67, 1358–1367.

Li, Y., Hoffmann, J., Li, Y., Stephano, F., Bruchhaus, I., Fink, C. and Roeder, T. (2016) Octopamine controls starvation resistance, life span and metabolic traits in *Drosophila*. Scientific Reports 19(6), 35359.

Li, Y., Tiedemann, L., Von Frieling, J., Nolte, S., El-Kholy, S., Stephano, F., Gelhaus, C., Bruchhaus, I., Fink, C. and Roeder, T. (2017) The role of monoaminergic neurotransmission for metabolic control in the fruit fly *Drosophila melanogaster*. Frontiers in Systems Neuroscience 11, 60.

Liao, M., Xiao, J-J., Zhou, L-J., Liu, Y., Wu, X-W., Hua, R-M., Wang, G-R. and Cao, H-Q. (2016). Insecticidal activity of *Melaleuca alternifolia* essential oil and RNA-seq analysis of *Sitophilus zeamais* transcriptome in response to oil fumigation. PLoS ONE 11, 12.

Ma, H., Huang, Q., Lai, X., Liu, J., Zhu, H., Zhou, Y., Deng, X. and Zhou, X. (2019). Pharmacological properties of the type 1 tyramine receptor in the Diamondback moth, *Plutella xylostella*. International Journal of Molecular Sciences 20, 2953.

Mannino, G., Abdi, G., Maffei, M.E. and Barbero, F. (2018). *Origanum vulgare* terpenoids modulate *Myrmica scabrinodis* brain biogenic amines and ant behaviour. PLoS ONE 13(12), e0209047.

Neckameyer, W.S. and Leal, S.M. (2017) Diverse functions of insect biogenic amines as neurotrasmitters, neuromodulators and neurohormones. In book: Hormones, Brain and Behaviour 2, 367–401.

Nishimura, T., Seto, A., Nakamura, K., Miyama, M., Nagao, T., Tamotsu, S., Yamaoka, R. and Ozaki, M. (2005). Experimental effects of appetitive and nonappetitive odors of feeding behavior in the blowfly, *Phormia regina*: a putative role for tyramine in appetite regulation. Journal of Neuroscience 25, 7507–7516.

Ohta, H. and Ozoe, Y. (2014). Molecular signalling, pharmacology, and physiology of octopamine and tyramine receptors as potential insect pest control targets. Advances in Insect Physiology 46, chapter two.

Orchard, I., Ramirez, J.M. and Lange, A.B. (1993). A multifunctional role for octopamine in Locust flight. Annual Review of Entomology 38, 227–249.

Park, C.G., Jang, M., Yoon, K.A. and Kim, J. (2016). Insecticidal and acetylcholinesterase inhibitory activities of Lamiaceae plant essential oils and their major components against *Drosophila suzukii* (Diptera: Drosophilidae). Industrial Crops and Products 89, 507–513.

Pauls, D., Blechschmidt, C., Frantzmann, F., Jundi, B. and Selcho, M. (2018). A comprehensive anatomical map of the peripheral octopaminergic/tyraminergic system of *Drosophila melanogaster*. Scientific Reports 8, 15314.

Ponton, F., Chapuis, M-P., Pernice, M., Sword, G.A. & Simpson, S.J. 2011. Evaluation of potential reference genes for reverse transcription-qPCR studies of physiological responses in *Drosophila melanogaster*. Journal of Insect Physiology 57: 840–850.

Price, D.N. and Berry, M.S. (2006). Comparison of effects of octopamine and insecticidal essential oils on activity in the nerve cord, foregut and dorsal unpaired median neurons of cockroaches. Journal of Insect Physiology 52, 309–319.

Priestley, C.M., Williamson, E.M., Wafford, K.A., Satelle and D.B. (2003). Thymol, a constituent of thyme essential oils, is a positive modulator of human GABA and a homo-oligosteric GABA receptor from *Drosophila melanogaster*. British Journal of Pharmacology 140, 1363–1372.

Regnault-Roger, C., Vincent, C. and Arnason J.T. (2012). Essential oils in insect control: low-risk products in a high-stakes world. Annual Review of Entomology 57, 405–424.

Rillich, J., Stevenson, P. and Pflueger, H. (2013). Flight and walking in locust-cholinergic co-activation, temporal coupling, and its modulation by biogenic amines. PLoS ONE 8, e62899.

Robertson, J.L., Jones, M.M., Olguin, E. and Alberts, B. (2017). Bioassays with arthropods. 3rd ed. Boca Raton, FL: CRC Press, Taylor & Francis Group.

Roeder, T. (2005). Tyramine and octopamine: ruling behaviour and metabolism. Annual Review of Entomology 50, 447–477.

Roeder, T. (2020). The control of metabolic traits by octopamine and tyramine in invertebrates. Journal of Experimental Biology 223, 194282.

Roeder, T., Seifert, M., Kähler, C. and Gewecke, M. (2003). Tyramine and octopamine: antagonist modulators of behavior and metabolism. Archives of Insect Biochemistry and Physiology 54, 1–13.

Rota-Stabelli, O., Blaxter, M. and Anfora, G. (2013). Drosophila suzukii. Current Biology 23, 8–9.

Saraswati, S., Fox, L.E., Soll, D.R. & Wu, C-F. (2004). Tyramine and octopamine have opposite effects on the locomotion of *Drosophila* larvae. Journal of Neurobiology 58(4), 425–41.

Saudou, F., Amlaiky, N., Plassat, J.L., Borrelli, E. and Hen, R. (1990). Cloning and characterization of a *Drosophila* tyramine receptor. The EMBO Journal 9(11), 3611–3617.

Schetelig, M.F., Lee, K.Z., Otto, S., Talmann, L., Stokl, J., Degenkolb, T., Vilcinskas, A. and Halitschke, R. (2017). Environmentally sustainable pest control options for *Drosophila suzukii*. Journal of Applied Entomology 142(1-2), 3–17.

Schützler, N., Girwert, C., Hügli, I., Mohana, G., Roignant, J-Y., Ryglewski, S and Duch, C. (2019). Tyramine action on motoneuron excitability and adaptable tyramine/octopamine ratios adjust *Drosophila* locomotion to nutritional state. PNAS 116, 3805–3810.

Velasco, B., Erclik, T., Shy, D., Sclafani, J., Lipshitz, H., McInnes, R & Hartenstein, V. (2006). Specification and development of the pars intercerebralis and pars lateralis, neuroendocrine command centers in the Drosophila brain. Developmental Biology 302(1), 309–323.

Walsh, D.B., Bolda, M.P., Goodhue, R.E., Dreves, A.J., Lee, J.C., Bruck, D.J., Walton, V.M., O’neal, S.D. and Zalom, F.G. (2011). *Drosophila suzukii* (Diptera: Drosophilidae): Invasive pest of ripening soft fruit expanding its geographic range and damage potential. Journal of Integrated Pest Management 1, 1–7.

Wu, S.F., Huang, J. and Ye, Y.Y. (2013). Molecular cloning and pharmacological characterisation of a tyramine receptor from the rice stem borer, *Chilo suppressalis* (Walker). Pest Management Science 69, 126–134.

Wu, S.F., Xu, G., Qi, Y.X., Xia, R.Y., Huang, J. and Ye, G.Y. (2014). Two splicing variants of a novel family of octopamine receptors with different signalling properties. Journal of Neurochemistry 129, 37–47.

Yatsenko, A.S., Marrone, A.K., Kucherenko, M.M and Shcherbata, H.R. (2014). Measurement of metabolic rate in *Drosophila* using respirometry. Journal of Visualized Experiments 24(88), e51681.

Zhai, Y., Lin, Q., Zhou, X., Zhang, X., Liu, T. & Yu, Y. (2014). Identification and validation of reference genes for quantitative real-time PCR in *Drosophila suzukii* (Diptera: Drosophilidae). PLoS ONE 9(9), e106800.

Zhang, Z., Yang, T., Zhang, Y., Wang, L and Xie Y. (2016). Fumigant toxicity of monoterpenes against fruitfly, *Drosophila melanogaster*. Industrial Crops and Products 81, 147–151.

